# *Akkermansia muciniphila* mediates antibiotic-induced impairment of female preference in mice

**DOI:** 10.1101/2025.09.19.677244

**Authors:** Maryana Morozova, Olga Snytnikova, Jelizaveta Andrejeva, Julia Popova, Kseniya Achasova, Yuri Tsentalovich, Elena Kozhevnikova

## Abstract

Antibiotic-induced disruptions of the gut microbiota have been linked to behavioral alterations, but the mechanisms underlying these effects remain poorly understood. Here, we demonstrate that combined vancomycin and rifampicin (VR) treatment in male C57BL/6 mice impairs female odor preference and reduces interaction time with female conspecifics in favor of males, without affecting general locomotion or anxiety-like behavior. VR-induced effects on the intestinal microbiome persisted for weeks after antibiotic withdrawal. Metabolomic profiling revealed brain ATP depletion upon vancomycin treatment, while intranasal administration of the electron transport chain uncoupler dinitrophenol (DNP) partially recapitulated antibiotics-induced social deficits. Microbiome analysis and functional tests identified *Akkermansia muciniphila* and short chain fatty acids acetate and propionate as key mediators of the behavioral phenotype. Strikingly, this VR-induced behavioral phenotype was reproducible in BALB/c mice, suggesting broad applicability. Our findings establish VR treatment as a model for microbiota-dependent social deficits, implicating mitochondrial metabolism and specific bacterial taxa in sexually dimorphic behaviors. These results highlight the need to consider neuroactive antibiotic effects in clinical and translational research.

## 1. Introduction

The gut microbiome has emerged as a critical regulator of brain function and behavior, with growing evidence linking microbial dysregulation to neurodevelopmental and psychiatric disorders [1], [2]. This relationship is mediated through the gut-brain axis, involving immune, metabolic, and neural pathways that influence social behavior, feeding, and stress responses [3]. In mice, social interactions ‒ such as mating, aggression, and territorial behavior ‒ are strongly dependent on olfactory cues [4]. The vomeronasal and main olfactory systems detect pheromones and social odors, which are processed in the hypothalamus and amygdala to drive sexually dimorphic behaviors [5]. For example, male mice exhibit innate preferences for female urinary odors [6], and demonstrate mating toward females and aggressive behaviors toward male intruders. While both mating and aggression have been shown to involve hypothalamic circuits related to estrogen-expressing neurons [7]. Disruptions in these systems, whether through genetic, hormonal, or environmental factors (including microbiota alterations), can profoundly alter social hierarchies and reproductive success [8].

The relationship between gut microbiota and social behavior has been increasingly recognized, with germ-free animal models showing significant alterations in social interaction patterns [9], [10]. However, less is known about how specific, targeted modifications of the microbiome might affect complex social behaviors, particularly those involving sex-specific preferences.

Recent studies have demonstrated that antibiotic-induced alterations of gut microbiota can lead to significant behavioral changes, including impaired social interaction and cognitive function [11], [12]. While the effects of broad-spectrum antibiotic cocktails on behavior have been reported, the specific contributions of individual antibiotics and their combinations remain poorly understood. This gap is particularly relevant given the widespread clinical use of antibiotics and their potential long-term neurological consequences [13].

Broad-spectrum antibiotics impair social odor discrimination and reduce affiliative interactions in mice [11], likely through disruptions in microbial metabolites like short-chain fatty acids that regulate neuroinflammation and blood-brain barrier integrity [14]. For instance, vancomycin, which targets Gram-positive bacteria, can deplete *Lactobacillus* and *Bacteroides* species that produce neuroactive GABA and serotonin precursors [15], [16], while rifampicin’s inhibition of mitochondrial RNA polymerase may compromise energy metabolism in neurons [17] Notably, antibiotic-treated mice exhibit reduced interest in female odors and impaired mate discrimination [18]. However, whether these effects arise from direct neuronal toxicity, microbiota shifts, or both remains unknown.

Recent evidence suggests that antibiotic-induced behavioral changes may be mediated through multiple pathways, including alterations in mitochondrial function, neurotransmitter metabolism, and immune signaling [19], [20], [21]. Of particular interest is the potential role of mitochondrial energy metabolism, as several antibiotics have been shown to affect ATP production and oxidative phosphorylation [17], [22]. The hypothalamus, a key regulator of social behavior and energy homeostasis, may be especially vulnerable to such metabolic perturbations [23], [24]. However, the specific mechanisms linking antibiotic-induced microbiota changes to hypothalamic function and social behavior remain unclear.

Previous work has shown that combinations of multiple antibiotics can impair odor preference and induce depressive-like behaviors in mice [12], [18], but these studies often used complex mixtures that make it difficult to identify specific microbial contributions. Our study addresses this limitation by focusing on the effects of two clinically relevant antibiotics ‒ vancomycin and rifampicin ‒ known for their distinct impacts on gut microbiota composition [17], [25]. The selection of vancomycin and rifampicin for this investigation was based on several considerations. Vancomycin, a glycopeptide antibiotic, primarily targets Gram-positive bacteria and has been shown to dramatically reduce microbial diversity [26]. Rifampicin, a bactericidal antibiotic effective against mycobacteria and some Gram-positive organisms, has demonstrated neuroprotective properties in some contexts while showing mitochondrial toxicity in others [17], [27]. This paradoxical nature of rifampicin’s effects on neural function makes it particularly interesting for studying microbiome-gut-brain interactions. Importantly, both antibiotics are commonly used in clinical practice, making their potential neurological effects highly relevant for human health [28].

This study was based on our previous work that combined vancomycin and rifampicin (VR) treatment to partially recapitulate the unique behavioral phenotype observed in a transgenic model of chronic colitis and dysbiosis [29]. We hypothesized that the effects of VR on the behavioral profile would be mediated through specific alterations in gut microbiota and subsequent impacts on brain metabolism. To test these hypotheses, we employed a comprehensive approach combining behavioral analyses, *16S rRNA* sequencing of gut microbiota, and brain metabolomic profiling. By combining behavioral analyses with multi-omics approaches, we aimed to establish causal links between specific microbial changes, brain metabolic alterations, and behavioral outcomes. We further investigated strain-dependent effects of VR and compared responses in C57BL/6 and BALB/c mice, two commonly used inbred strains with known differences in social behavior and stress responses. Our integrated approach revealed key metabolic and microbial alterations that contribute to female preference in male mice.

## 2. Materials and Methods

### 2.1. Animals

The experiments involving animals were conducted at the Scientific Research Institute of Neurosciences and Medicine (SRINM) and at the Institute of Molecular and Cellular Biology (IMCB). Experimental procedures adhered to the regulations by Russian law, including Good Laboratory Practice as outlined in the Directive #267 from June 19, 2003 by the Ministry of Health of the Russian Federation. The procedures adhered to the institutional ethical committee’s guidelines, and the European Convention for the protection of vertebrate animals. The animal study protocol was approved by the Institutional Ethics Committee of The Institute of Molecular and Cellular Biology of 06.10.2021. All animals were tested for pathogens quarterly in accordance with the recommendations by the Federation of European Laboratory Animal Science Associations (FELASA) [30]. The research utilized the C57BL/6JNskrc (C57BL/6 further on) and BALB/cNskrc (BALB/c further on) strains, our in-house derivatives of C57BL/6J and BALB/cJ mouse strains respectively.

The study involved adult male mice 8-12 weeks of age housed in open cages (with a size of 318 × 202 × 135 mm, #CP-3, ABTex, Russia) with birch sawdust as bedding and paper cups as environmental enrichment. A 12-h light/dark cycle was maintained [12:00 (OFF): 00:00 (ON)] during housing and behavioral tests. Food (BioPro, Novosibirsk, Russia) and water were provided *ad libitum*. During pregnancy and before weaning, mice received dog chow pellets as protein supplements (Dry dog food for adult, small breed, “Opti Balance” with chicken, Purina Pro Plan, USA). Sexually experienced age-adjusted BALB/c (for C57BL/6 residents) and C57BL/6 (for BALB/c residents) males and estrus females were used as a source of bedding odor and as intruders. The choice of the intruder strain was based on easily recognizable fur color as compared to the test animals and its successful use in our previous studies [31], [32].

For the social odor preference tests and the two-intruders tests, sexually experienced males were used, as sexual experience strongly affects the outcome of such tests [31], [33], [34]. Sexual experience in test males was induced by co-housing with sexually mature females for a week (one male and one female per cage), after which the test males were single-housed throughout the experiment. Two or more biological replicates were performed for the most social odor and two-intruders tests in most groups. All animals were randomly allocated to the experimental groups.

Test animals were euthanized by craniocervical dislocation. Intestinal and brain samples were collected and frozen in liquid nitrogen for further analyses.

### 2.2. Antibiotic treatment

Antibiotics were provided in drinking water for two weeks following the scheme: ampicillin + clavulanic acid (A) was supplemented on day 0, vancomycin (V) was supplemented at day 0, rifampicin (R) was added at day7 either to water or to vancomycin solution (VR). The concentrations were: vancomycin ‒ 0.5 g/L, rifampicin ‒ 0.075 g/L, amoxicillin + clavulanic acid ‒ 0.45 g/L + 0.15 g/L, respectively. At the time of experimental tests, the animals continued to drink antibiotic solutions. Control (C) mice received drinking water. Feces for microbiome analyses were collected directly from animals in three consecutive days starting the day of the first behavioral test. Several biological replicates were performed for some groups as follows:

***Experiment 1***, C57BL/6: Control (n = 20), Vancomycin (n = 10), Rifampicin (n = 11), Vancomycin + Rifampicin (n = 11), Amoxicillin (n = 10).

***Experiment 2***, C57BL/6: Control (n = 10), Rifampicin (n = 10), Vancomycin + Rifampicin (n = 11), Amoxicillin (n = 10).

***Experiment 3***, C57BL/6: Control (n = 9), Vancomycin (n = 10), Vancomycin + Rifampicin (n = 10).

***Experiment 4***, BALB/c: Control (n = 11), Vancomycin + Rifampicin (n = 12) (as shown in Figure 5H).

***Withdrawal experiment:*** antibiotics were supplemented following the scheme described above. On day 17, antibiotics were withdrawn, and behavioral tests were performed 7, 14, and 25 days after withdrawal as shown in the scheme (Figure 5A). The control group was a separate group of animals (n = 12), while the VR-treated group (n = 12) was used in all withdrawal experiments.

### 2.3. Akkermansia muciniphila treatment

*Akkermansia muciniphila* (64013T, CCUG, Sweden) was cultivated on 10 cm Petri dishes with chocolate agar (#938500183805215, BioMedia, Russia) under anaerobic conditions at 37°C for 72 h in an anaerostatic jar (2.5 L, Thermo Fisher Scientific, USA) with gas-generating sachets (#AN0025A, Thermo Fisher Scientific, USA). Anaerobic conditions were confirmed by Oxoid Resazurin Anaerobic Indicator (#BR0055B, Thermo Fisher Scientific, USA). After cultivation, the bacterial colonies were washed from Petry dishes with sterile 10% glycerol in 1x PBS, aliquoted and stored at -70°C. The CFU count was estimated by a serial dilution method with subsequent culture on Petri dishes.

C57BL/6 mice were subjected to the intragastric gavage with *Akkermansia muciniphila* culture in 1x PBS with glycerol (10%) at 10^9^CFU in 200 µl. The gavage was performed every 3 days for 2 weeks (Figure 4C), and while the behavioral tests were performed. *Akkermansia muciniphila* supplementation proceeded until the end of the behavioral experiments on test-free days. Control group received 1x PBS with 10% glycerol gavage.

*Akkermansia muciniphila* supplementation was performed in two biological replicates. The first replicate involved control (n = 10) and *Akkermansia muciniphila* treated males (n = 9), while the replicate contained n = 9 males in both groups (n = 18-19 males per group in total).

### 2.4. Intranasal dinitrophenol (DNP) treatment

Dinitrophenol (DNP) (#6-09-1883-86, PJSC “Khimprom”, Russia) intranasal injection was performed in two independent biological replicates using two dosages of 2.5 and 4 mg/ml. DNP was resolved in 1x PBS and injected 40 min prior to the test in the volume of 15 µl (37.5 and 60 µg per animal in two nostrils, 7.5 µl per nostril). PBS intranasal injection served as control. There was no statistical difference between the two dosages, thus we combined the results in the same graphs (Figure 2F-G). Both replicates contained n = 10 males in both groups (n = 20 males per group in total). DNP concentration was chosen based on previously published data [35].

### 2.7. Short-chain fatty acids treatment

Short-chain fatty acids treatment was performed in two biological replicates.

***Experiment 1***: sodium acetate (Am-O602, Helicon, Russia) and calcium propionate (NX1011, Lianyungang Tongyuan Biotechnology, China) (both referred to as SCFA further on) were dissolved in drinking water at concentrations of 50 mM each and supplied to C57BL/6 mice (n = 10), while a control group of mice received regular water (n = 10). After two weeks of treatment, the behavioral tests were performed as shown in the schematic (Figure 4I). The treatment was proceeded throughout the behavioral experiments.

***Experiment 2***: SCFA were prepared and supplemented as described above (n = 10), while the control group received sodium chloride (50 mM) and calcium chloride (50 mM) in the drinking water (n = 9) to control for the effect of sodium and calcium ions on behavior. The concentration of SCFA was chosen according to previous studies and based on the approximated physiological levels [36], [37]. The data from both experiments was combined and analyzed together.

### 2.7. Spectroscopy

The metabolomic composition of the brain samples (n = 8 per group) from the control, V and VR-treated groups was studied at the Center of Collective Use “Mass spectrometric investigations” of the SB RAS in Novosibirsk, Russia, using high-resolution 1H NMR spectroscopy. Brains were cut symmetrically in the sagittal plane and flash frozen in liquid nitrogen. Half of each weighed brain samples was then homogenized in a cold (-20°C) mixture of water, methanol, and chloroform at a 1:2:2 volume ratio, using 1.6 mL per 150 mg of tissue to extract water-soluble metabolites for quantitative NMR metabolomic analysis. After homogenization, the mixtures were vortexed, cooled on ice, and incubated at -20°C, followed by centrifugation to pellet proteins. The top aqueous layer was collected, dried by a vacuum concentrator, and then dissolved in 600 μl of deuterated phosphate buffer (0.01 M, pH = 7.4) containing 6 µM sodium 4,4-dimethyl-4-silapentane-1-sulfonate (DSS) as a standard. The detailed sample preparation protocol was previously published in [38], [39].

The obtained extracts were analyzed on a Bruker BioSpin AVANCE III HD 700 MHz NMR spectrometer. The 1H NMR spectra were recorded with multiple scans at a consistent 25°C using a 90-degree detection pulse, pre-saturation for the water signal, and adequate relaxation time between scans. Manual spectral correction such as phasing and baseline, as well as signal processing and peak integration were performed using MestReNova V.12 (Mestrelab Research S.L., Spain) software. Metabolite identification was cross-referenced against the Human Metabolome Database (http://www.hmdb.ca, accessed on January 12, 2024) and previous metabolomic profiling studies, and validated by adding reference compounds as needed [29], [38], [39]. The absolute concentration of metabolites was determined by the integrating the peak area of the metabolite relative to the DSS standard.

### 2.8. Behavioral tests

Behavioral tests were performed in sexually experienced male mice, which were single-housed in open cages (318 × 202 × 135 mm, #CP-3, 3W, Russia). The interval between tests was 2-3 days. The behavioral tests were performed in the following order: open field test, non-social odor discrimination and habituation, female odor discrimination test, odor preference test, sociability test, female contact test, resident-intruder test, and the two-intruders test (Figure 1A).

**Figure 1.**
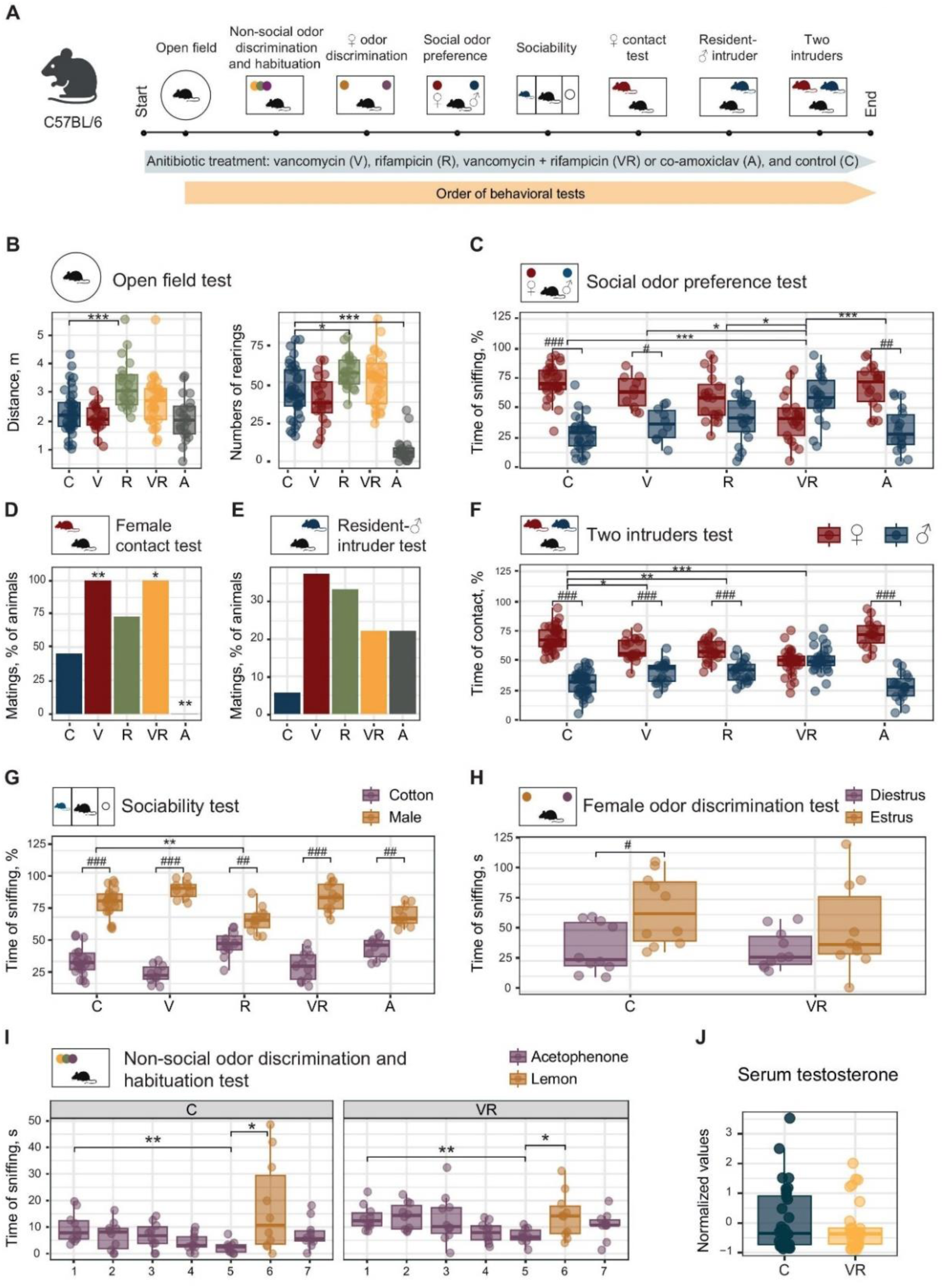
Combined rifampicin and vancomycin treatment impair odor preference and social preference toward females. **A.** Schematic of the antibiotic treatment experiment with C57BL/6 male mice. **B.** Open field test (*n* = 20-38). One-way ANOVA with Tukey post-test, *\* p* < 0.05, *\*\*\* p* < 0.001. **C.** Social odor preference test shown as a percent of time spend sniffing social odors (*n* = 10-21). One-way ANOVA with Tukey post-test, *\* p* < 0.05, *\*\*\* p* < 0.001. Within group comparisons (between stimuli) were performed with paired Student’s t-test with BH adjustment, *# p* < 0.05, *## p* < 0.01, *### p* < 0.001. **D.** Female intruder test (*n* = 10-20). Pairwise comparisons were made using Exact Fisher test with BH adjustment. **E.** Male intruder test (*n* = 8-18). **F.** Two-intruders test. Social communication is shown as a percent of time spend with both intruders (*n* = 18-39). One-way ANOVA with Tukey post-test, *\* p* < 0.05, *\*\* p* < 0.01, *\*\*\* p* < 0.001. Within group comparisons (between stimuli) were performed with paired Student’s t-test with BH adjustment, *### p* < 0.001. **G.** Sociability test. Social communication is shown as a percent of time spend with both stimuli (*n* = 10-20). One-way ANOVA with Tukey post-test, *\*\* p* < 0.01. Within group comparisons (between stimuli) were performed with paired Student’s t-test with BH adjustment, *## p* < 0.01, *### p* < 0.001. **H.** Female odor preference test is shown as time sniffing estrous and diestrus females’ urine (s) (*n* = 10 in both groups). Repeated measures ANOVA followed by paired Student’s t-test with Benjamini-Hochberg (BH) adjustment, *# p* < 0.05. **I.** Odor discrimination and habituation. Data is shown as time sniffing a specimen (s) (*n* = 10 in both groups). Repeated measures ANOVA followed by paired Student’s t-test with BH adjustment for planned comparisons, *\* p* < 0.05, *\*\* p* < 0.01. **J.** Testosterone measurements in blood serum (*n* = 29-32). Wilcoxon sum rank test. Box plots represent median ± IQR.

**Figure 2.**
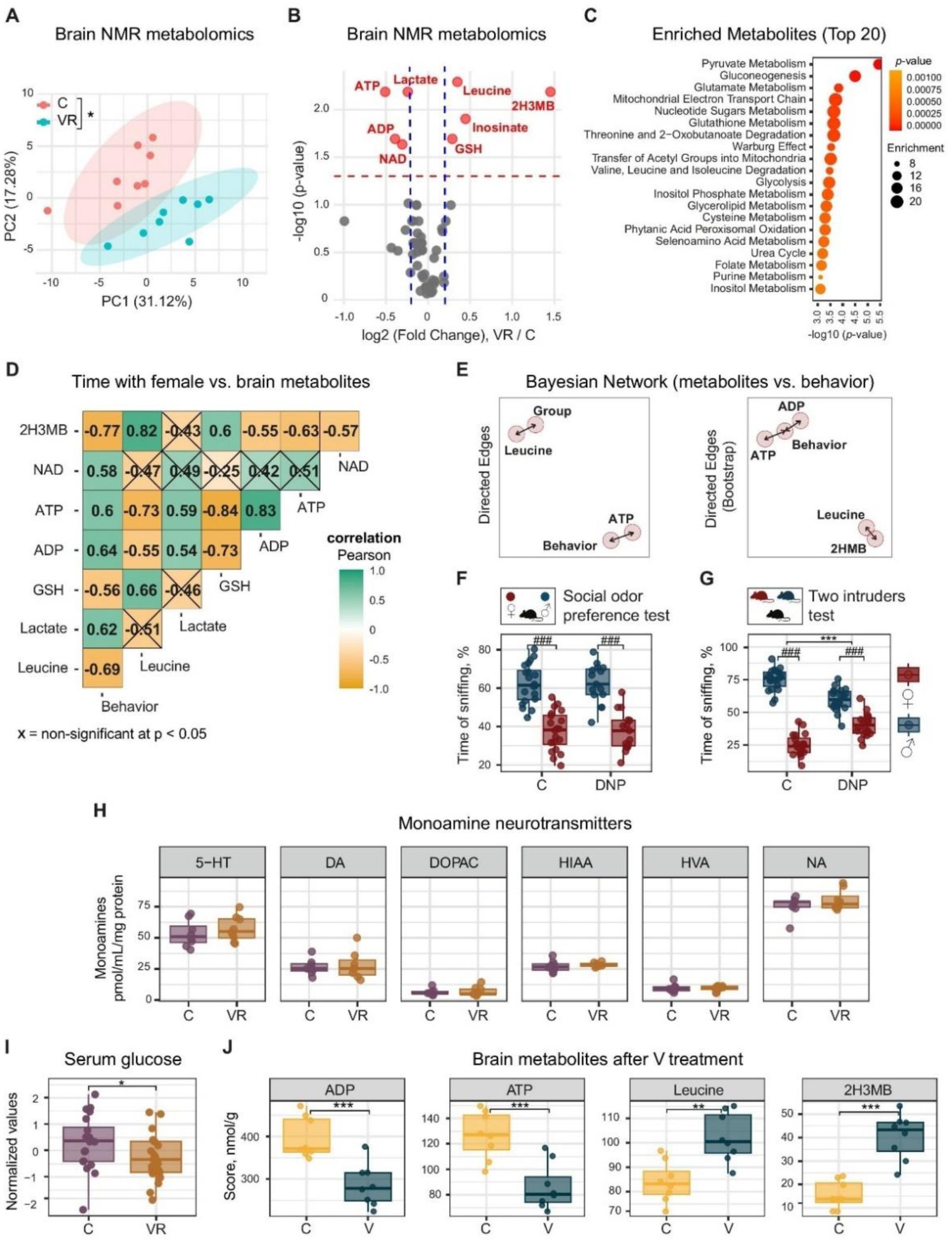
Brain metabolism after combined rifampicin and vancomycin treatment. **A.** PC analysis of brain metabolic profiles (*n* = 8 in each group). Student’s t-test, *p* < 0.05. **B.** Volcano plot of mean metabolite concentrations vs. *p*-values of mean differences (*n* = 8 in each group). Student’s t-test with BH adjustment. Metabolites with absolute value of foldchange > 1.2 (blue dashed lines) and *p* < 0.05 (red dashed line) are shown in red. **C.** A list of enriched metabolite sets for significantly different metabolites from Figure 2B. The graph shows enrichment ratio (circle size) and *p*-value (dot color) as assessed by the Fisher’s exact test. **D.** Pearson correlation matrix of the selected metabolites with the female contact time in the Two-intruders test (*n* = 16). Non-significant correlations (*p* ≥ 0.05 after BH adjustment) with correlation coefficient < 0.05 are crossed out. **E.** Bayesian network build upon selected metabolites and time with female before and after bootstrapping (*n* = 16, 100 permutations). Moderately stable edges were allowed. ATP <-> “behavior”: *p* = 0.005, ADP <-> “behavior”: *p* = 0.005, 2H3MB <-> leucine: *p* = 0.0003, Pearson correlation tests at *α* = 0.05. **F.** Social odor preference test after intranasal DNP treatment (*n* = 15-20). Circles and triangles represent different doses. Within group comparisons (between stimuli) were performed with paired Student’s t-test with BH adjustment, *### p* < 0.001. **G.** Two-intruders test after intranasal DNP treatment (*n* = 20 for both groups). Circles and triangles represent different doses. Between group comparisons (within one stimulus) were made Student’s t-test for independent samples with BH adjustment, *\*\*\* p* < 0.001. Within group comparisons (between stimuli) were performed with paired Student’s t-test for dependent samples with BH adjustment, *### p* < 0.001. **H.** Monoamines measured in hypothalamic samples normalized to protein content (*n* = 6-8). **I.** Serum glucose concentration showed as standardized values (*n* = 20-21). Wilcoxon Rank Sum test, *\* p* < 0.05. **J.** Brain metabolites after V treatment (*n* = 8 for both groups). Student’s t-test with BH adjustment, *\*\* p* < 0.01, *\*\*\* p* < 0.001. Box plots represent median ± IQR. 5-HT – serotonin, DA – dopamine, DOPAC – 3,4-Dihydroxyphenylacetic acid, HIAA – 5-Hydroxyindoleacetic acid, HVA – 4-hydroxy-3-methoxy-phenylacetic acid, NA – noradrenaline, 2H3MB – 2-hydroxy-3-methyl-butyrate.

Behavioral tests after antibiotic treatment involving C57BL/6 mice were performed in three independent experiments. The first experiment included all five animal groups and was conducted in two parallel cohorts. The first cohort consisted of two antibiotic-treated groups and a control group, tested on the same day. The second cohort, comprising the remaining two treatment groups along with an additional control group, was tested the following day. This setup enabled efficient testing of multiple animals within a short timeframe while ensuring adequate rest periods between experiments. The following tests were performed during the first experiment: open field, social odor preference, sociability, female contact test, resident-intruder, and the two-intruders test. The second experiment included open field, non-social odor discrimination and habituation, female odor discrimination, social odor preference, and two-intruders test. The third experiment included open field and two-intruders test. The results of all available biological replicates were combined per treatment group and shown in Figure 1.

Behavioral tests after antibiotic treatment involving BALB/c mice were performed as one biological replicate and included open field and two-intruders test (Figure 5H).

In the withdrawal experiment, there were two groups of C57BL/6 male mice: a control group received no treatment and was tested in the social odor preference and two-intruders tests. Another group of males received VR treatment and was tested in the social odor preference and two-intruders tests. After that, VR was withdrawn and was animals were tested in the social odor preference test until the odor preference to females recovered. After that, the two-intruders test was performed (Figure 5A).

DNP-treated animals were subjected to the social odor preference and two-intruders tests (Figure 2F-G).

Mice after *Akkermansia muciniphila* gavage were tested in the open field, odor preference, social odor preference, and the two-intruders tests (Figure 4C).

Animals that received SCFA in drinking water were tested in odor preference, social odor preference, and the two-intruders tests (Figure 4I).

Some animals were excluded from the analysis based on the following criteria: total odor/intruder investigation times were < 20 s, which correspond to the average time an animal spends in the random area of the cage [34]. Males showing highly aggressive behavior toward a female intruder, which resulted in wounding, were also excluded from analysis. Some animals were excluded from the open field test analysis because their total distance traveled fell outside three standard deviations from the mean. The number of animals per each test is shown in figure legends.

#### 2.8.1. Open field test

Open field test measures motor and exploratory activity, and can also reflect general anxiety in a test animal. For this experiment, a mouse was introduced into the center of a clear-walled 40 × 40 cm square arena under white light. Central square of 20 × 20 cm in size was considered as the center. An overhead video camera captured the mouse’s locomotion, whereas a separate camera positioned laterally tracked its vertical movements for 6 min. Total distance, time spent in the center of the arena, the number of rearings and the number of entries in center were analyzed using the Ethovision XT10 (Noldus, Netherlands) software.

#### 2.8.2. Odor preference test

Dry dog food pellets (“Opti Balance” with chicken, Purina Pro Plan, USA) were placed in tea infusers (#469.568.00, Ikea, Sweden) as a source of the attractive non-social odor for 5 min. Glass beads resembling dog food in shape and size were used as sham controls. As food deprivation is often used to stimulate preference toward food odor [40], test animals were food-deprived 12 h before the test. Test mice were weighed before and after food deprivation, ensuring weight loss does not exceed 10-15%. Active investigation involving characteristic movements of the nose and whiskers was recorded as “sniffing” behavior and was evaluated manually. The results are given as “time spent sniffing a specimen” / “total sniffing time” and expressed as percentages.

#### 2.8.3. Non-social odor discrimination, olfactory habituation, and olfactory memory

The home cage lid was substituted by a plastic cover with a hole to place cotton swabs soaked in odor samples. 0.01% acetophenone (#42163, Sigma-Aldrich, Missouri, United States) and 0.01% lemon flavor (Lemon, Golden Garden, Spain) were used for olfactory habituation and discrimination of non-social odors. First, acetophenone was presented five consecutive times for 3 min each with a one-minute interval. Then the lemon odor was presented for 3 min. After 60 min, acetophenone was presented for another 3 min.

#### 2.8.4. Female odor discrimination test

This experimental setting allows to evaluate the discrimination between two female odors. The test relies on male mice’s innate preference for estrous female urine over diestrus urine. For this test, the same setting was used. First, diestrus female urine was presented for 2 min followed by the estrus female urine after a 5 min interval.

#### 2.8.5. Social odor preference test

The social odor preference test investigates animal’s interest to female and male bedding odors without direct interaction with the odor source. Bedding samples from both female and male BALB/c mice were placed into the test animal’s home cage in two tea infusers (#469.568.00, Ikea, Sweden) for 5 min. BALB/c animals were kept in their cages unchanged for a week before the test in order to soil the bedding. Active investigation involving characteristic movements of the nose and whiskers was recorded as “sniffing” behavior and was evaluated manually. The results are given as “time spent sniffing a specimen” / “total sniffing time” and expressed as percentages.

#### 2.8.6. Sociability test

This test evaluates autistic-like behavior and demonstrates preference towards animals rather than unanimated objects and was conducted as described previously [29]. Briefly, a rectangular arena (60 × 40 cm) divided into three equal chambers (20 × 40 cm) with 6 × 6 cm openings under red light was used as a test environment. An animal was placed in the central chamber and allowed to move freely for 5 min for acclimation. An unfamiliar BALB/c male mouse and a white cotton ball of comparable size were placed in the either side of the chamber under wire cups. Time spent in each chamber was analyzed automatically using Ethovision XT10 (Noldus, Netherlands) software. Social preference was calculated as “time spent in the “animal” chamber” / “time spent in the “cotton” chamber. Choosing a chamber to place a BALB/c mouse was random.

#### 2.8.7. Contact tests

These tests were used to investigate social interest and social interactions like sniffing, aggression and courtship behavior. Three-month-old sexually experienced BALB/c animals were used as intruders. Either a female, a male or both intruders were placed in the test animal’s home cage for 15 min [29], and social behavior was recorded using video camera. Attacks and matings were counted manually during the test by two observers and averaged out. Indicators of social interest, such as the frequency and time of sniffing were calculated in video by the same observers and subsequently averaged. The female intruder test was performed at first followed by the male intruder test, and the two intruders test was performed last. The time of sniffing (nasal and genital), grooming, chasing, and following were considered as the time of contact and expressed as a percentage of the total contact time with a male and a female intruder.

### 2.9. Metagenomic analysis

For the metagenomic analysis, fecal samples were collected in Inhibitex buffer, and total DNA was extracted using QIAamp DNA Stool Mini Kit (#51604, QIAGEN, Germany) according to manufacturer’s instructions (n = 6-7 per group). Metagenomic analysis was performed by a commercial service at Novogene (https://en.novogene.com, accessed on January 20, 2025) as follows. The 16S rRNA V3-V4 region was used for barcoded amplification using Phusion High-Fidelity PCR Master Mix (#M0531S, New England Biolabs, UK), and PCR-products were mixed in equal ratios. The mixture was purified with Qiagen Gel Extraction Kit (#28704, QIAGEN, Germany). The libraries were generated with NEBNext Ultra DNA Library Prep Kit for Illumina (#E7645S, New England Biolabs, UK) sequencing and analyzed by the Illumina platform. Data analyses were performed using Uparse and QIIME (Version 1.7.0) software. Sequences with ≥ 97% similarity were assigned to the same operational taxonomic units (OTUs). Principal component analysis (PCA) preceded cluster analysis to reduce the dimension of the original variables using the FactoMineR package and ggplot2 package in R software (Version 2.15.3). Principal Coordinates Analysis (PCoA) was performed using top 70 genera, identified in at least one of the test groups, top two PCs in terms of variance coverage were used for data plotting and identification of significant differences among the groups.

### 2.10. Testosterone and glucose measurements

Blood samples (n = 29-32) were collected from mice (three experiments) using an orbital sinus puncture and centrifuged at 3,000 rpm for 15 min at room temperature. Then, 20 μL of blood sera were used to measure testosterone by ELISA using the commercial kit according to the manufacturer’s protocol (#X-3972, Vector-Best, Novosibirsk, Russia). Serum samples (n = 20-21) from two VR treatment experiments were used for glucose measurement using ELISA kit according to the manufacturer’s instructions (#B-8056, Vector-Best, Novosibirsk, Russia). Glucose concentrations were standardized to account for the effect of separate measurements and shown as z-standardized values. Blood samples from *Akkermansia muciniphila* gavage (n = 8-9) are shown as actual concentration values.

### 2.11. Monoamine measurements

Monoamines were extracted from hypothalamic samples (n = 8 in both groups). Tissue fragments were homogenized in 0.1 M HClO₄. Homogenization was performed using a Q125 ultrasonic homogenizer (Qsonica, USA), followed by centrifugation (10 min, 4°C; 14,000 *g*). Monoamine measurements were conducted using high pressure liquid chromatography with electrochemical detection (Shimadzu, Japan). Monoamine concentrations were calibrated using commercial standards and normalized to total protein in a sample measured with Bradford assay.

### 2.12. Statistical analyses

The data are presented as box plots with overlaying individual data points and 1.5 interquartile ranges (IQR), or bar graphs for percentages using ggplot2 package of R software (Version 4.4.1). To test the normality of distributions, the Shapiro-Wilk test was employed. Data that followed a normal distribution was evaluated using analysis of variance (ANOVA) following the Student’s t-test for independent or dependent samples. Non-normally distributed data were analyzed with Kruskal-Wallis test with Wilcoxon sum rank test as post hoc. The percentage of animals that attacked or mated in the two-intruders test was analyzed with the Fisher exact test. Benjamini-Hochberg (BH) procedure was applied to all experiments with more than two groups to adjust for multiple comparisons.

For metagenomic analyses of antibiotic-treated groups, taxa comparisons were performed using one-way ANOVA with Tukey HSD test. Metagenomic composition after antibiotic withdrawal was analyzed with one-way PERMANOVA with 999 permutations followed by Wilcoxon sum rank test as post hoc because the control group was independent of the other three measurements performed in the VR-treated group repeatedly. PCA and PCoA were assessed by PREMANOVA with 999 permutations. Linear Discriminant Analysis (LDA) Effect Size (LEfSe) results are represented as cladograms where only taxa with LDA score > 4 and adjusted *p*-value < 0.05 are shown.

Metabolomic data were analyzed using z-standardized values. Metabolite comparisons between groups were performed using Student’s t-test with BH adjustment. Enriched metabolites were analyzed with MetaboAnalyst 6.0 (https://www.metaboanalyst.ca/, accessed on January 16, 2025).

Correlation analyses were performed with either Pearson or Spearman test with BH adjustment depending on data distribution.

Bayesian network was constructed using significantly enriched metabolites and the time spend in contact with a female in the two-intruders test. We used a constraint-based Bayesian network approach with the PC (Peter-Clark) algorithm with the following parameters: conditional independence testing via Pearson’s correlation, a significance threshold of *α* = 0.05. Edge stability was assessed through 100 bootstrap iterations, retaining edges appearing in > 35% of resampled datasets. Only significant edges (*p* < 0.05, Pearson’s correlation) are shown. The analysis was performed in Python (Version 3.12.7), the PC algorithm was implemented using pgmpy package[41].

A *p*-value of less than 0.05 was considered indicative of statistical significance.

## 3. Results

### 3.1. Combined rifampicin and vancomycin treatment impairs odor preference and social preference toward females

To assess the effect of antibiotics on social odor discrimination and preference as well as their impact on social behavior with male and female conspecifics, we established five experimental groups of male mice each receiving either vancomycin (V), rifampicin (R), vancomycin + rifampicin (VR), amoxicillin + clavulanic acid (A) in water, and a control (C) group supplemented with water only (Figure 1A). Test C57BL/6 mice received antibiotics for 14 days followed by the behavioral tests in the order shown in Figure 1A. The treatment was proceeded throughout the behavioral experiments. Open filed test revealed that none of the treatments inhibited overall locomotor and horizontal investigatory activity (Figure 1B). However, rifampicin treatment increased the distance travelled in the open filed and the number of rearings during the test, indicating increased activity and investigation. Amoxicillin treatment strongly inhibited vertical activity (rearings), without affecting total distance (Figure 1B). The result of the open field test revealed no major inhibitory effect of antibiotics on general locomotion and investigation, and no sickness behavior was observed.

In order to evaluate the impact of antibiotics on social odor perception, the social odor preference test was performed (Figure 1C). It revealed that there was an effect of treatment on female odor preference. It was substantially reduced in VR group as compared to the control, the R and V treatments alone, and A group. This result suggests that the combined action of these antibiotics affects odor preference rather than the antibiotics *per se*. Control mice preferred sniffing female odor as well as amoxicillin -and vancomycin-treated males (Figure 1C). At the same time, there was no female preference in R and VR experimental groups indicating that some effect on social odor preference was achieved by the addition of rifampicin alone.

When tested with either a male or a female intruder in the separate resident-intruder tests, V- and VR-treated mice demonstrated more matings with female intruders (Figure 1D). There was an effect of the treatment on matings and interaction time with a female. Vancomycin alone and combined with rifampicin substantially increased the number of matings with female intruders.

At the same time, there were no differences in the resident-intruder test with a male intruder (Figure 1E).

To directly compare interest in males and females, we performed the two-intruders test, that combines both female and male animals in the home cage of a male resident. This experiment revealed that vancomycin, rifampicin and their combination reduced female preference in test males in comparison to the control group. However, the lack of preference to the female within group was only observed in the VR treatment, but not in other groups (Figure 1F). These results suggest that V and R treatments have an effect on male’s preference to a female, but their combination enhances this effect.

To estimate the overall sociability of mice upon antibiotic treatment, we performed a three-chamber test evaluating preference to a live animal rather than an unanimated object (Figure 1G). There was an effect of treatment on social preference to the conspecific, however, only rifampicin affected social preference to a living object as compared to the control. In all groups, mice preferred communicating with an animal rather than mock (Figure 1G).

Given that combined vancomycin and rifampicin treatment demonstrated the strongest effect on both female odor and social preference, we tested the effect of the combined VR treatment on social and non-social odor discrimination. The urine from either estrous or diestrous females was used as an odor stimulus for the female odor discrimination test (Figure 1H). It is expected that wild-type male mice prefer sniffing estrous female’s urine, and their interest increases when estrous odor is presented after the diestrous sample. Indeed, control male mice spend more time sniffing estrous sample, whereas VR-treated males demonstrated only tendency to preference, which was not statistically significant (Figure 1H). To evaluate whether this effect could be attributed to the general impairment of olfaction in the VR-treated mice, we further tested the odor discrimination and habituation to non-social odors. We found that both control and VR groups habituate to the odor of acetophenone and remember it, while discriminating lemon odor presented after a set of trials with acetophenone (Figure 1I). This result suggests that the general olfaction is not affected upon VR treatment.

As sexual behavior and aggression as well as perception and processing of female- and male-derived odors are regulated by androgens, we measured testosterone in the VR-treated males. There was no significant difference in serum testosterone between the control and VR-treated males (Figure 1J), suggesting the mechanism other that androgen regulation via VR.

### 3.2. Vancomycin impairs brain mitochondrial metabolism

The described above behavioral experiments identified VR treatment as a potent regulator of social behavior in mice. To further assess the effect of these antibiotics on the central metabolism and neurotransmitter biosynthesis, we evaluated brain metabolome by NMR spectroscopy and monoamine profile using liquid chromatography.

Brain metabolome profiling of the control and VR groups identified 55 small compounds, including intermediary metabolites, amino acids, sugars, nucleic acids, and others. Based on these profiles, principal component analysis (PCA) revealed that VR treatment significantly shifted brain metabolism as compared to the control (Figure 2A). Of these metabolites, 8 significantly changed at least 1.2-fold (absolute value) (Figure 2B). Among these were leucine, 2-hydroxy-3-methyl-butyrate (2H3MB), inosine, glutathione (GSH), that were upregulated upon VR intake (Figure 2B). At the same time, ATP, ADP, NAD, and Lactate were downregulated in the brain of VR males (Figure 2B). The enrichment analysis of metabolites based on the metabolic pathway demonstrated that among the most prevalent are mitochondrial processes, including electron transport chain, pyruvate and glutamate metabolism, and others (Figure 2C). An apparent change in ATP, ADP, NAD and Lactate was indicative of VR interference with mitochondrial metabolism, however its relation with social behavior was unclear.

Thus, we assessed a possible association between the identified metabolites and female preference in the two-intruders test (shown as “behavior”). Correlation analysis revealed that most listed metabolites correlate with the percentage of time a male resident spend with a female intruder (Figure 2D). Likewise, almost all metabolites correlated (positively or negatively) with each other, suggesting their metabolic relation or relation to the experimental group (Figure 2D). Given this data, it is impossible to identify any key metabolite over its counterparts.

In order to propose possible casual relationships and further associations between the metabolic compounds in the brain and female preference, we tested directed probabilistic relationships between metabolites and social behavior using a constraint-based Bayesian network. For this analysis, we chose metabolites whose correlation with behavior was over 0.7 to minimize the number of predictors and increase the robustness of the model (Figure 2D). These are 2H3MB, leucine, ATP, and ADP, which were combined with the percentage of time spent with a female intruder in the two-intruders test as another variable (“behavior”). The Bayesian network structure identified significant relations between ATP, ADP, and the behavioral parameter, and between 2H3MB and leucine (Figure 2E). This approach did not gain enough power to result in the directional casual relationships, however, it revealed that ATP is more likely to be related to the behavioral phenotype observed in the two-intruders test. The identified edges were confirmed using bootstrapping approach (Figure 2E, right panel).

To evaluate biological relevance of the identified above relation, we tested the effect of ATP downregulation on odor and social preference. We intranasally applied an electron transport chain uncoupler dinitrophenol (DNP) to male mice 40 min prior to the tests. An intranasal treatment was chosen to localize the effect of DNP to the olfactory system and the downstream neuronal connections, while reducing the systemic effect of the uncoupler. Social odor preference test revealed no effect of DNP treatment on female odor preference (Figure 2F). In the two-intruders test, DNP-treated males also preferred females, however, female preference was significantly weaker than that in the control group (Figure 2G). This result suggests that ATP is indeed involved in regulation of female and male preference, and might contribute to the phenotype observed upon the antibiotic treatment.

The NMR analysis also identified key metabolites involved in multiple types of behavior such as glutamate and gamma-amino-butyric acid, but no significant difference was found for these compounds. Likewise, monoamine neurotransmitters measured by liquid chromatography in hypothalamic samples of VR-treated males did not differ between the two groups (Figure 2H). Given this result, so far, APT remained our sole candidate key metabolite involved in social preference toward females.

Next, we testes whether reduced ATP levels originate from the lowered blood glucose levels reported in some antibiotic treatment experiments [42]. In agreement with previous studies, a reduction of blood glucose was observed after VR treatment in our experimental set-up (Figure 2I). Thus, a possible explanation of the reduced mitochondrial metabolism is the systemic loss of glucose induced by VR.

Finally, we tested whether vancomycin treatment itself could affect ATP levels in the brain given that some reports suggest its direct inhibitory effect on mitochondrial activity [43]. We performed NMR analysis of the brain samples from mice treated with vancomycin alone. As can be seen, ATP increase can be fully attributed to the effect of vancomycin, as the ATP level in the V group was also reduced (Figure 2J). In agreement with this, the effect of the intranasal DNP treatment on the two-intruders test outcome largely mirrors that of the vancomycin treatment alone (compare Figures 1F and 2G). Therefore, we conclude that behavioral phenotype of the VR treatment at least in part originates from the effect of vancomycin on ATP production in the brain.

### 3.3. Microbial signatures in the intestine after rifampicin and vancomycin treatment

However, it can be seen that only a minor portion of the behavioral phenotype can be explained by the ATP reduction, thus, there must be other factors that influence behavior upon VR treatment. Thus, we investigated the microbiome of mice treated with the antibiotics to understand the impact of their antibacterial effects on male behavior.

To evaluate the effect of each antibiotic and their combination on the bacterial composition in the intestine, we analyzed stool samples from the test and control animals by the 16S rRNA gene sequencing. A stronger effect on the overall microbiota composition was achieved by vancomycin as compared to rifampicin as can be seen by weighted unifrac PCoA (Figure 3A). However, the addition of rifampicin (in the VR group) attenuated the effect of vancomycin on the bacterial profile so that V and VR compose distinct groups in terms of microbiome composition (Figure 3A). In addition to this, the biodiversity of the microbiome was significantly reduced in the V and VR groups as revealed by the Shannon’s and Simpsons’s diversity indexes (Figure 3B-C), while there was no significant difference between the control and R groups regarding these parameters.

**Figure 3.**
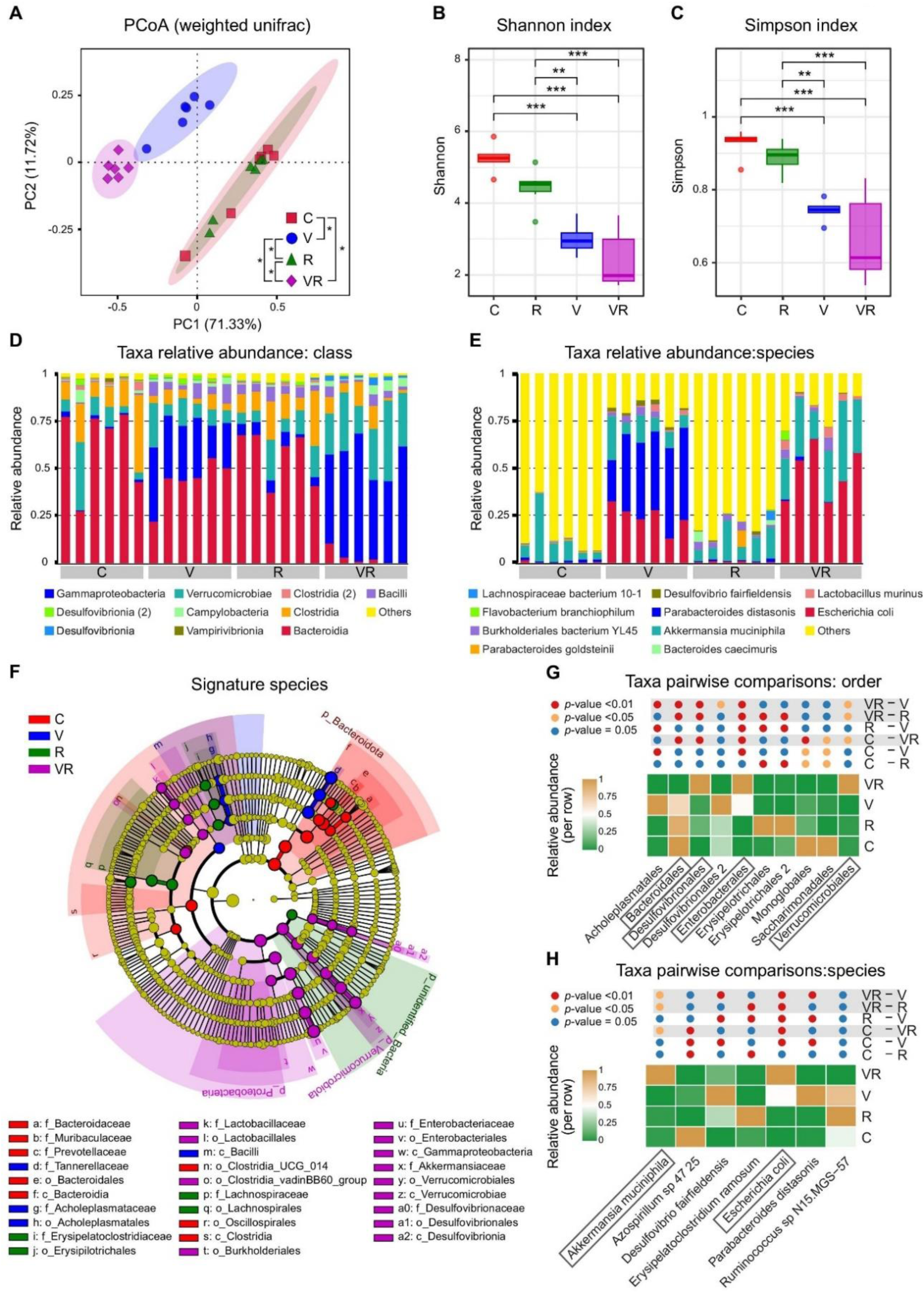
Analysis of the microbial profiles after antibiotic treatment. (*n* = 6 in each group). **A.** PCoA analysis of the intergroup differences. Two-way PERMANOVA with pairwise comparisons, *\*\* p* < 0.05 (BH adjusted). **B-C.** Shannon and Simpson indices. Box plots represent median ± IQR, Two-way ANOVA with Tukey post-test; *\* p* < 0.01, *\*\* p* < 0.01, *\*\*\* p* < 0.001). **D-E.** Bar plots of normalized bacterial abundances at the class (D) and species level (E). **F.** Cladogram representing signature species for the four treatment groups. LEfSe analysis, only signatures with LDA score above 4 and *p* < 0.05 are shown. **G-H.** Intergroup comparisons at the order (G) and species (H) level. Heatmaps represent relative abundance per taxonomic group. Only taxa with significant V and R factors interaction are shown (Two-way ANOVA). The result of Tukey post-test is color coded as shown in the legend. The pairs of comparisons where significant differences were expected are highlighted in grey. Significantly different taxa in all expected pairs of comparisons are shown in boxes.

**Figure 4.**
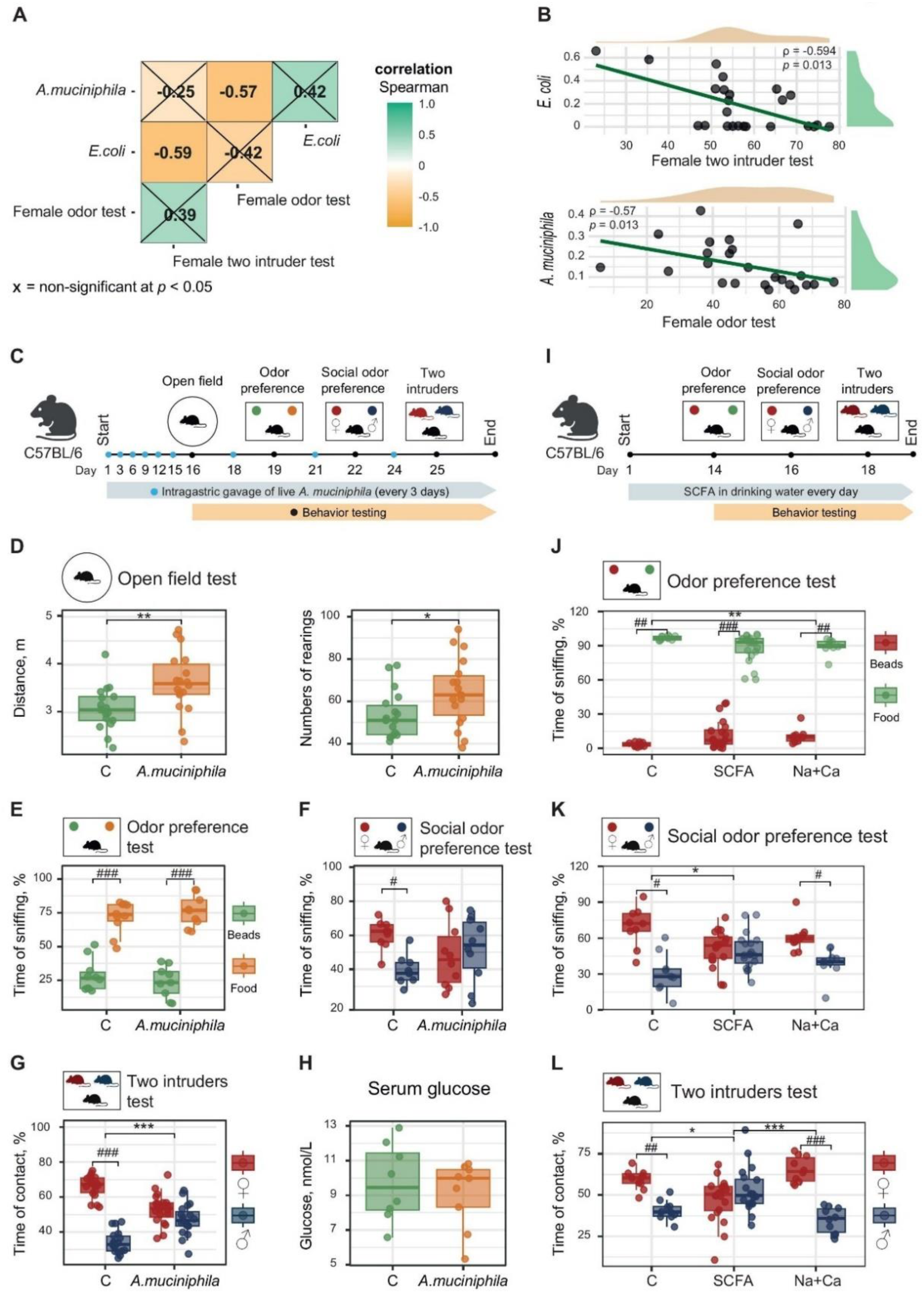
*Akkermansia muciniphila* and SCFA impair female odor and social preference. **A.** Correlation matrix between female preference in the social odor test and in the two-intruders test expressed as the percentage of social interaction time with normalized bacterial count of *Akkermansia muciniphila* and *Escherichia coli* (*n* = 24). Spearman test with BH adjustment, non-significant correlations at *p*<0.05 are crossed out. **B.** Individual correlation plots for significant pairs (*n* = 24). Spearman test with BH adjustment. **C.** Schematic of *Akkermansia muciniphila* gavage experiment. **D.** Open filed test after A. muciniphila gavage (*n* = 17-18 in each group). Distance: Student’s t-test, *\*\* p* < 0.01; rearings: Wilcoxon Rank Sum test, *\* p* < 0.05. **E.** Odor preference test (*n* = 9-10). **F.** Social odor preference test after *Akkermansia muciniphila* gavage (*n* = 9-10). **G.** Two-intruders test after *Akkermansia muciniphila* gavage (*n* = 18-19). **H.** Glucose after *Akkermansia* muciniphila gavage (n = 8-9). **I.** Schematic of SCFA supplementation experiment. **J.** Odor preference test after SCFA supplementation (*n* = 9-20). **K.** Social odor preference test (*n* = 9-20). **L.** Two-intruders test after SCFA supplementation (*n* = 9-19). In **E-G** and **J-L**: Between group comparisons (within one stimulus) were made using Student’s t-test for independent samples with BH adjustment, *\*\* p* < 0.01, *\*\*\* p* < 0.001. Within group comparisons (between two stimuli) were made using Student’s t-test for dependent samples with BH adjustment, *## p* < 0.01, *### p* < 0.001. Box plots represent median ± IQR. SCFA – sodium acetate and calcium propionate, Na+Ca – sodium and calcium chloride.

**Figure 5.**
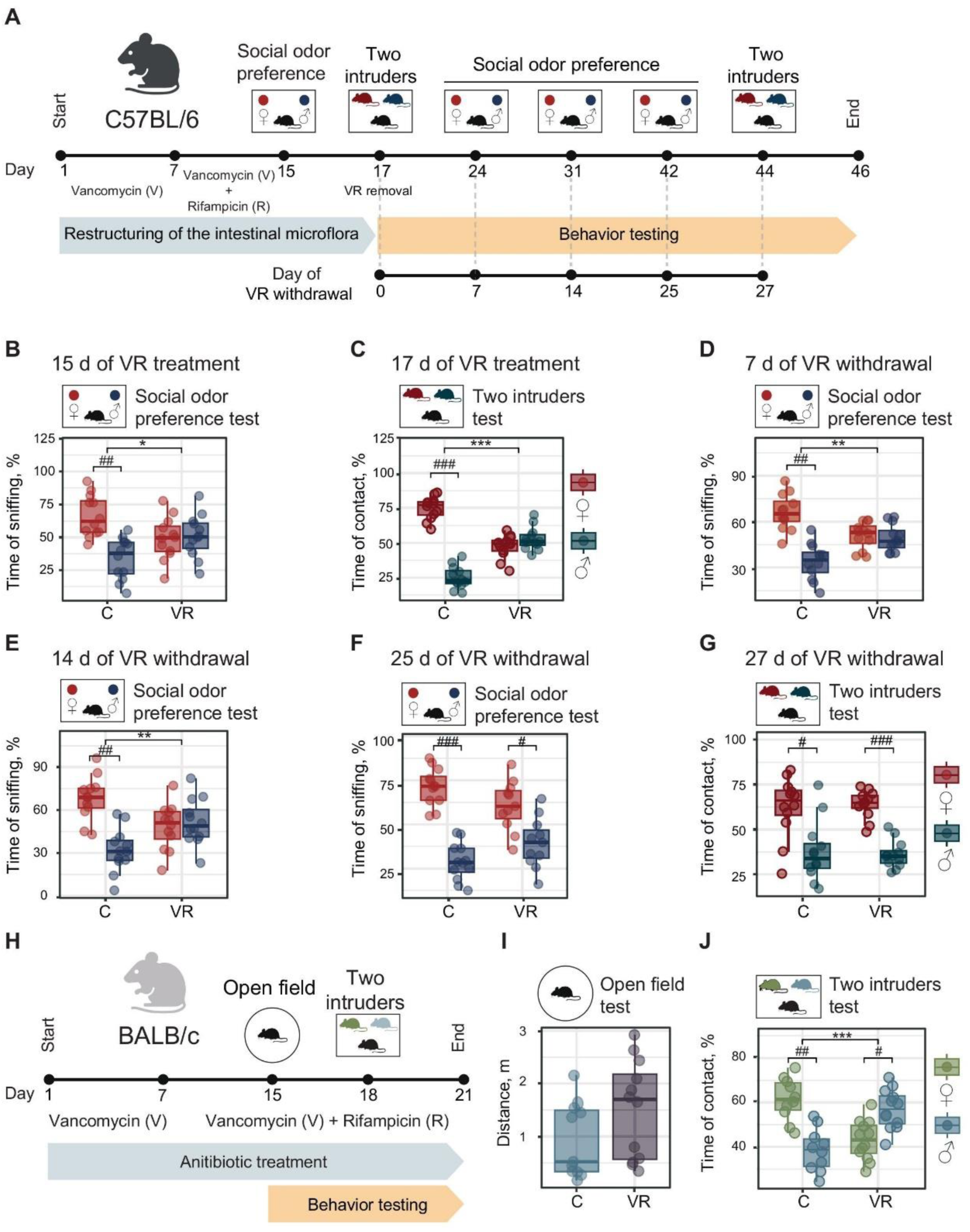
Combined vancomycin and rifampicin treatment and withdrawal as a model of social preference deficit. Vancomycin (V), rifampicin (R), vancomycin + rifampicin (VR) in water, and a control (C) group supplemented with water only. **A.** Timeline of the VR treatment and withdrawal in C57BL/6 mice. **B.** Social odor preference test after 2 weeks of VR treatment (day 15) (*n* = 12 in each group). **C.** Two-intruders test after 2 weeks of VR treatment (day 17) (*n* = 12 in each group). **D.** Social odor preference test after 7 days of VR withdrawal (day 24) (*n* = 12 in each group). **E.** Social odor preference test after 14 days of VR withdrawal (day 31) (*n* = 12 in each group). **F.** Social odor preference test after 25 days of VR withdrawal (day 42) (*n* = 11-12). **G.** Two-intruders test after 27 days of VR withdrawal (day 44) (*n* = 12 in each group). **H.** Timeline of the VR-treatment experiment in BALB/c mice. **I.** Open field test after two weeks of VR treatment (*n* = 11-12). **J.** Two-intruders test after 2 weeks of VR treatment (day 18) (*n* = 10-12). Between group comparisons (within one stimulus) were made using Student’s t-test for independent samples with BH adjustment, *\* p* < 0.01, *\*\* p* < 0.01, *\*\*\* p* < 0.001. Within group comparisons (between two stimuli) were made using Student’s t-test for dependent samples with BH adjustment, *# p* < 0.01, *## p* < 0.01, *### p* < 0.001. Box plots represent median ± IQR.

We then investigated the abundance of the bacterial taxa in the experimental groups and found that all antibiotic treatments introduced some changes in the microbiome composition. Again, the strongest effect was achieved in the V and VR groups at all taxonomic levels (the data for the classes and species are shown in Figure 3D-E). Notably, the apparent difference was also observed between the V and VR groups at all levels, with a strong decrease of *Bacteroida* (red), and an increase of *Verrucomicrobiae* (turquoise) classes in the VR group as compared to the V group (Figure 3D).

At the species level, *Escherichia coli* (red) was strongly elevated in both V and VR groups as compared to the control, while an increase of *Parabacteroides diastonis* (blue) evoked by vancomycin was eliminated by the addition of rifampicin (Figure 3E). Although the biodiversity was as low in the VR as in the V group, an increase in *Escherichia coli* and *Akkermansia muciniphila* (turquoise) compensated for the lack of *Parabacteroides diastonis* under the combined antibiotic treatment (Figure 3E).

LEfSe (linear discriminant analysis (LDA) Effect Size, [44]) analysis revealed biomarker taxa associated with V vs control, and V vs VR groups (Figure 3F). Interestingly, these biomarker groups partially overlap. Comparing C and VR reveals key features that might explain the behavioral differences resulting from the antibiotic treatment, while comparing V with VR helps to narrow down the taxa potentially responsible for the phenotypes found in the VR group only. Thus, *Verrucomicrobiales*, *Enterobacterales*, *Lactobacillaes*, and *Desulfovibrionales* are the most promising biomarker taxa associated specifically with VR treatment. These bacterial species might be involved in the behavioral outcome found in this animal group (Figure 3F).

At the order level, there was an effect of treatment on the 10 abundant species (species were retained for analysis only if they exceeded 0.1% relative abundance in all samples within at least one experimental group) (Figure 3G). Again, only bacteria that distinguish VR from all control, V, and R groups can be considered as putatively associated with the behavioral phenotype as it was only observed upon the combined antibiotic treatment. We then searched for taxa that were specifically changed in the VR groups as compared to control, V, and R groups (Figure 3G, highlighted in grey). Within 10 taxa, *Bacteroidales*, *Desulfovibrionales*, *Enterobacterales*, and *Verrucomicrobiales* satisfied this condition and thus could be involved in regulation of social behavior (Figure 3G, shown in boxes).

Within species, only *Akkermansia muciniphila* and *Escherichia coli* satisfy the considerations elaborated above, however, *Escherichia coli* also upregulates in the V group as compared to the control although to the lesser extent (Figure 3H). Given the result of microbiome analysis, *Akkermansia muciniphila* and *Escherichia coli* remain the potential species to be tested in further experiments (Figure 3H, shown in boxes).

### 3.4. Akkermansia muciniphila gavage reduces female odor and intruder preference in C57BL/6 mice

Next, we tested whether our key features *Akkermansia muciniphila* and *Escherichia coli* correlate with behavioral parameters and can serve as candidate species that might be associated with the impaired social behavior with male and female animals and their odors. We performed Spearman correlation analysis of *Akkermansia muciniphila* and *Escherichia coli* abundances with the percentage of time spend by the test males either sniffing females’ odor samples or contacting females in the two-intruders test (Figure 4A-B). The result shows that *Akkermansia muciniphila* negatively correlates with the female odor sniffing, while *Escherichia coli* correlates with the time spend in direct contact with a female intruder. Importantly, the abundancies of both bacteria also have positive correlation suggesting some relation between these bacterial species (Figure 4A). At the same time, preference to female odor and preference to a female intruder have no significant correlation, which has been observed and reported before (Figure 4A, [31]).

Given that *Akkermansia muciniphila’s* intestinal abundance correlates with the lack of interest toward the female odor, we tested the effect of its oral gavage on the odor and intruder preference. The odor preference was chosen as a parameter that is most likely unrelated to the ATP production. Isolated culture of *Akkermansia muciniphila* was delivered by oral gavage to C57BL/6 adult males. Then the open field, the food odor preference, the social odor preference, and the two-intruders tests were performed as shown on the schematic (Figure 4C). The open field test suggested that *Akkermansia muciniphila* induced horizontal and vertical exploratory activity resulting in the increased distance and number of rearings (Figure 4D). This agrees with the elevated vertical activity upon the VR treatment.

The intragastric gavage of cultured *Akkermansia muciniphila* did not affect food odor discrimination, which was shown using the corresponding test, suggesting that this bacterium does not affect general olfaction (Figure 4E). At the same time, *Akkermansia muciniphila* reduced female odor preference in the social odor preference test (Figure 4F). However, there was no significant difference between the control and treated males regarding female odor preference. Likewise, test males also exhibited less interest to the female intruder than the control males in the two-intruders test (Figure 4G) and did not prefer interacting with female intruders to males (Figure 4G). These results suggest that indeed, the abundance of the intestinal *Akkermansia muciniphila* is related to male-male and male-female behavior. In contrast to VR treatment, there was no significant effect on blood glucose after *Akkermansia muciniphila* gavage (Figure 4H).

Given that *Akkermansia muciniphila* is a major acetate and propionate producer, we evaluated the effect of acetic and propionic acids (“SCFA” further on), two short chain fatty acids produced by *Akkermansia muciniphila* on the female odor preference and social interaction (schematic of the experiment is shown in Figure 4I). Since SCFA were administered as calcium and sodium salts, the control group received either drinking water or a mixture of the corresponding chloride salts. Like *Akkermansia muciniphila* gavage, SCFA did not affect food odor preference in males (Figure 4J). At the same time, SCFA supplementation decreased female odor preference in the social odor preference test (Figure 4K), and contact time with a female in the two-intruders test (Figure 4L). Thus, both *Akkermansia muciniphila* and SCFA induce similar behavioral phenotypes, suggesting the involvement of bacterial metabolism in odor preference and social behavior.

### 3.5. Combined vancomycin and rifampicin treatment as a model of social preference deficit

The next experiment was developed to analyze the duration of the antibiotics’ effect on male social odor preference and social communication with males and females. The antibiotics were supplemented in water for two weeks followed by regular water. The social tests were set after two weeks of treatment (no withdrawal), and 7, 14, and 25 days post withdrawal (Figure 5A). After VR removal, only social odor test was performed until the female odor preference restored. After that, the odor test was complemented by the two-intruders test (Figure 5A).

At the point of VR treatment, there was a reduction in sniffing time of female sample (Figure 5B) and a reduction of interest toward females in the contact test (Figure 5C). On the days 7 and 14 days of withdrawal, there was no restoration of interest to female odor, and the time of contact with female samples was significantly lower in the withdrawal group than in the control (Figure 5D-E). The odor preference restored only by the 25^th^ day after withdrawal (Figure 5F). Likewise, interaction time with female intruders in the two-intruders test also increased (Figure 5G). It is worth noting that the adverse behavioral effects of antibiotics persist beyond the initial treatment period.

We also tested whether the combination of the two antibiotics induce similar effects on male and female preference in another inbred mouse strain. Here we used BALB/c mice (Figure 5H-J), which are widely demanded for *in vivo* research. No adverse effect of VR treatment on motor activity or general anxiety was observed in this mouse strain in the open field test, which aligns well with our findings in C57BL/6 mice (Figure 5I). VR treatment resulted in the similar phenotype in the two-intruders test in BALB/c males with even stronger preference to male mice (Figure 5J). Only 20% of the test males preferred female intruders in contrast to about 80% of untreated BALB/c mice. This result suggest that VR treatment protocol can be translated to other mouse models and strains, which will allow creating precise *in vivo* tools for behavioral studies and translational research.

### 3.6. Combined vancomycin and rifampicin treatment induces long-term microbial shifts in the intestine

Finally, we investigated the prolonged effect of VR treatment on the intestinal microbiome aiming to evaluate the duration of its antimicrobial effect and search for other bacterial modifiers of social behavior. Interestingly, 16S sequencing revealed that the major changes introduced by VR resolved neither 2 weeks, nor 3.5 weeks after antibiotics withdrawal (Figure 6A). PCoA analysis revealed that the overall microbiome profiles from the three treatment groups (VR, 14 days withdrawal, and 25 days withdrawal) differed from the control group, but were similar to each other (Figure 6A). Likewise, alfa diversity reflected in Shannon and Simpson indices was reduced after the VR treatment, and did not restore after 14 or 25 days of withdrawal (Figure 6B-C). This indicates that antibiotics not only decreased species richness, but also increased dominance of fewer taxa. There was no overall recovery in the biodiversity after withdrawal (Figure 6B), however, more species became dominant after 14 days (Figure 6C). This effect was not significant after 25 days of withdrawal, probably, due to the high variance in this group (Figure 6C).

**Figure 6.**
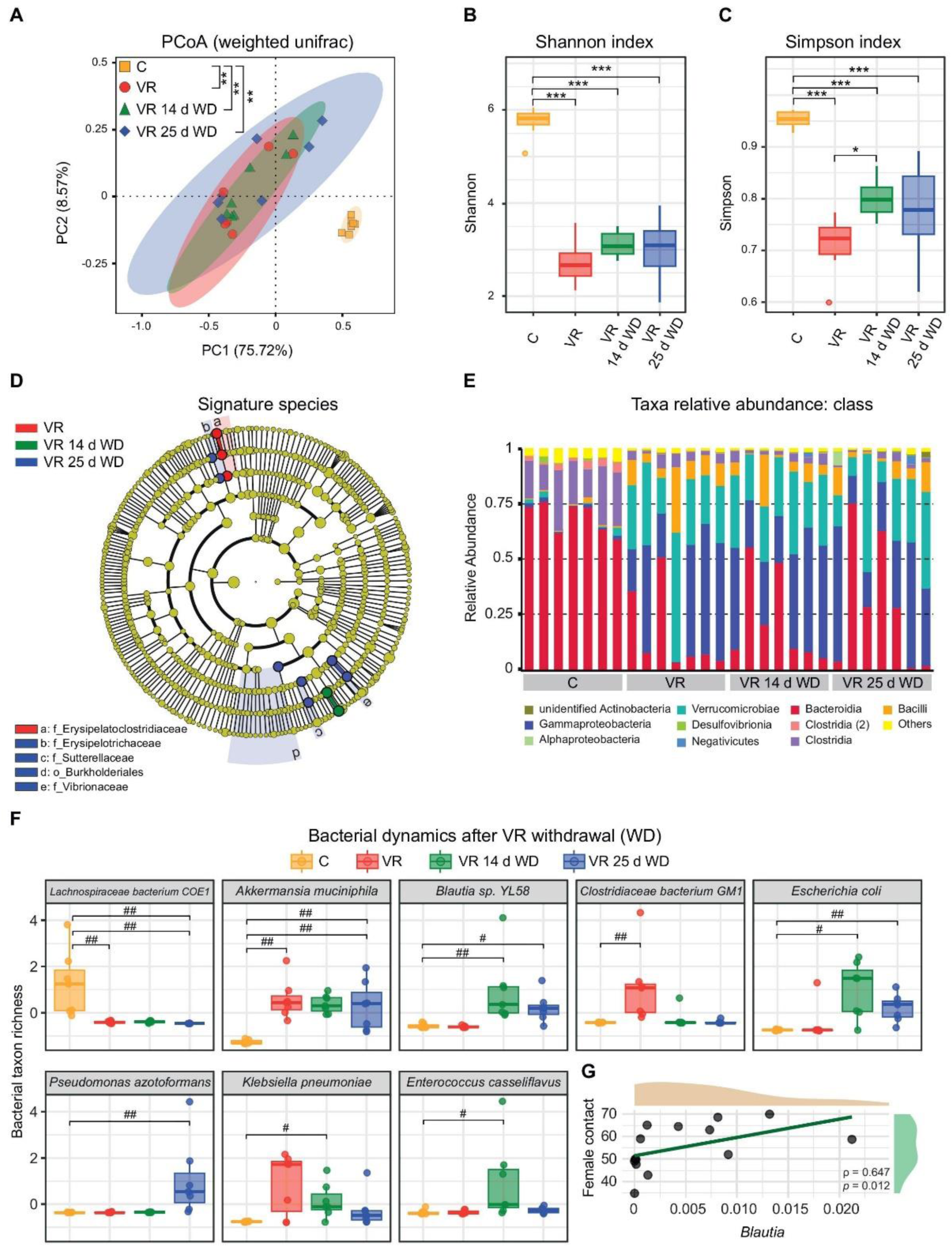
Combined vancomycin and rifampicin (VR) treatment induces long-term impairment of the intestinal microbiota. Vancomycin (V), rifampicin (R), vancomycin + rifampicin (VR) in water, and a control (C) group supplemented with water only. **A.** PCoA of the microbial features before and after VR withdrawal (*n* = 7 in each group). Two-way PERMANOVA with pairwise comparisons, *\*\* p* < 0.05 (BH adjusted). **B-C.** Shannon and Simpson indices (*n* = 7 in each group). Box plots represent median ± IQR, Two-way ANOVA with Tukey post-test; *\* p* < 0.01, *\*\* p* < 0.01, *\*\*\* p* < 0.001. **D.** Cladogram representing signature species for the four treatment groups (*n* = 7 for both groups). LEfSe analysis, only signatures with LDA score above 4 and *p* < 0.05 are shown. **E.** Bar plot of normalized bacterial abundances at the class level. **F.** Intergroup comparisons at the species level (*n* = 7 for all groups). Bar plots represent relative abundance per taxonomic group. Only taxa with the significant effect of the group are shown (One-way PERMANOVA with 999 permutations). Comparisons between groups were made using paired (when comparing treatment groups, *\* p* < 0.05) or unpaired (when comparing to the control group, *# p* < 0.01, *## p* < 0.01, *### p* < 0.001) Wilcoxon Rank Sum test. **G.** Spearman correlation between *Blautia* and female contact time in the two-intruders test 7 days and 25 days after withdrawal (*n* = 14). Box plots represent median ± IQR.

We further analyzed potential signature species that could discriminate VR treatment group from the withdrawal groups using LDA, but did not find many associated features except for *Sutterelaceae* and *Vibrionaceae* (Figure 6D). The bar plot of major species comparison between the four groups also shows that there was no apparent difference between dominant taxa between VT treatment and withdrawal groups. Interestingly, *Akkermansia muciniphila* (*Verrucomicrobiae*) was strongly upregulated after VR treatment in comparison to the control as was seen in the first experiment.

We next searched for taxa that change abundance under the four conditions using dispersion analysis and searched for those, where the factor of the “group” had significant influence at the species level. We found 8 bacterial species that were affected in this experiment including *Akkermansia muciniphila* and *Escherichia coli*. However, the latter was not upregulated after two weeks of VR administration, and only upregulated after withdrawal (Figure 6F). This result does not support the findings in the first experiment, where this bacterium propagated during the antibiotic treatment (Figure 3H).

Notably, *Akkermansia muciniphila* was overrepresented after 25 days of VR removal and was not different between the treatment and the withdrawal groups (Figure 6F). Given the behavioral pattern in the odor preference test, *Akkermansia muciniphila* profile does not fully explain social deficits that originate from VR exposure. Thus, other bacteria must be involved in the regulation of social odor preference in our model. Based on the post hoc comparisons, candidate bacteria include *Blautia sp. YL58* and *Pseudomonas azotoformans*, as their levels gradually change upon withdrawal (Figure 6F).

*Klebsiella pneumonia* upregulate after VR treatment and gradually decrease after withdrawal, however, these changes are not statistically significant (Figure 6E). Also, *Klebsiella pneumonia* is a pathogenic bacterium that proliferate upon antibiotic treatment. However, it was absent from the mouse cohorts in the first experiment, thus, it is unlikely to cause the phenotype.

*Pseudomonas azotoformans*, in turn, propagates specifically 25 days after withdrawal, and could be a good candidate to balance out the effect of *Akkermansia muciniphila* upregulation (Figure 6E). However, there is limited data on the effects of this microorganism in the intestine as it is an environmental species. The remaining candidate is *Blautia sp. YL58* as it increases with the withdrawal. Although there is no significant difference in *Blautia* between the 14^th^ and 25^th^ days of withdrawal, the overall trend suggests that this bacterial species might be associated with social behavior. We performed correlation analysis between *Blautia* abundance vs time of sniffing female odor and contact time in the two-intruders test, and revealed moderate correlation between the contact time with a female and *Blautia* (Figure 6G). Correlation with sniffing time was insignificant. Thus, *Blautia* species might be another potential candidate to regulate sexually dimorphic behaviors upon the withdrawal of antibiotics.

## 4. Discussion

The findings of this study demonstrate that combined VR treatment in male mice leads to significant and specific impairments in social behavior, particularly in interactions with female conspecifics. These effects manifest as reduced preference for female odors in social odor tests and diminished interaction time with female intruders.

Very similar results were reported by Salia and co-authors, who combined five antibiotics: ampicillin, neomycin, vancomycin, erythromycin, and gentamicin to demonstrate the role of intestinal microbiota in odor sensing [18]. The authors show that this antibiotic combination impairs female odor preference in males. However, other authors has shown previously that a combination of ampicillin, vancomycin, ciprofloxacin, imipenem and metronidazole might induce depressive-like behaviors in mice [12]. The behavioral changes observed in our study were not secondary to general locomotor or exploratory deficits potentially attributed to depression or anxiety, as open field tests confirmed normal activity levels across all treatment groups. This difference between our and other studies might arise from the choice of antibiotics used in different experiments. Our data support this idea by demonstrating that the combination of vancomycin and rifampicin produces unique behavioral signatures distinct from either antibiotic alone or ampicillin.

At the neural level, our metabolomic analysis revealed significant disturbances in brain energy metabolism following VR treatment. The observed reduction in ATP, ADP, NAD, and lactate levels suggest impaired mitochondrial function, similar to previous findings revealing the effect of antibiotics on brain metabolism [45]. Some antibiotic cocktails have been shown to affect systemic glucose metabolism, which may partially explain the downregulated energy metabolism in the brain [46]. Our findings agree with these studies: the VR-treatment resulted in lower blood glucose levels (Figure 2I). Likewise, some authors report that antibiotics affect brain neuro-transmitter metabolism including monoamines and tryptophan. These changes have been associated with social deficits [12]. However, we did not find significant change in neurotransmitters levels except for ATP (Figure 2B, H).

The Bayesian network analysis specifically linked the observed energy deficit to impaired social behavior, providing a potential mechanism for the behavioral changes in the VR model. At the same time, it is possible that the change in mitochondrial function and ATP levels are induced by vancomycin itself, because it has potential to affect mitochondrial activity [47]. The effect of vancomycin on brain mitochondrial metabolism is further confirmed by the NMR metabolomic profiling that shows similar changes in metabolites after vancomycin treatment (Figure 2J). Mitochondrial respiratory chain uncoupling demonstrated that ATP downregulation only partially explains the decrease of female preference, and does not affect female odor recognition (Figure 2F-G). This result is consistent with the effect observed for vancomycin treatment alone where test animals tend to lose interest toward females, but to a lesser extent than those treated with VR (Figure 1F). Thus, it is likely that the observed phenotype is at least partially explained by the reduction in energy metabolism evoked by vancomycin.

The role of rifampicin is more complex: there are contradicting studies regarding its role in mitochondrial metabolism [48], [49], [50], [51]. Some authors show that it protects mitochondria and promotes ATP production, while others demonstrate its toxic effect and ATP decrease. Here we see that rifampicin also tend to decrease time with a female (Figure 1F), so it is impossible to completely exclude the direct effect of both antibiotics on the two-intruders test in the VR group. It might be different for the odor preference test, where neither rifampicin, not vancomycin alone have any statistically significant effect (Figure 1C). The persistence of the behavioral effects for 3.5 weeks after antibiotic withdrawal also suggests against the direct effects of antibiotics on this type of behavior. It is more likely that a long-term microbial community disruption affected social preference. This finding extends the observations of Leclercq et al. (2017) who reported prolonged behavioral effects following early-life antibiotic exposure [11].[9] Likewise, our previous studies also showed a crucial role of microbiota in regulation of social behavior and social preference in the contexts of intestinal inflammation-induced dysbiosis [29], [52].

The antibiotics used in this study exert distinct effects on the intestinal microbial species. Vancomycin, which primarily targets Gram-positive bacteria, reduces obligate anaerobes such as *Clostridium* clusters IV and XIVa (including *Roseburia* and *Faecalibacterium prausnitzii*), *Bifidobacterium spp.*, and *Lactobacillus spp.*, while promoting the expansion of vancomycin-resistant enterococci (VRE) and *Proteobacteria*. Rifampicin suppresses both Gram-positive (*Bacteroides*, *Prevotella*) and Gram-negative (*Escherichia*, *Klebsiella*) taxa, often leading to a decline in *Firmicutes* and an increase in opportunistic pathogens such as *Enterococcus spp*. Ampicillin significantly depletes *Bacteroidetes* (particularly *Bacteroides spp*.) and certain *Firmicutes* (e.g., *Lactobacillus* and *Enterococcus*), while favoring the proliferation of ampicillin-resistant *Proteobacteria* (e.g., *Escherichia coli*) and fungi such as *Candida spp*. However, ampicillin treatment had no effect on female odor preference and time in contact with a female intruder.

The combination of vancomycin and rifampicin was designed in our previous study to specifically promote the outgrowth of *Akkermansia muciniphila* while inhibiting *Escherichia coli* [29]. Vancomycin treatment resulted in the most dramatic reduction of microbial diversity, consistent with previous reports [25]. However, the addition of rifampicin created a unique microbial signature characterized by enrichment of *Verrucomicrobiales*.

The experiment on antibiotic withdrawal does not support our initial finding that *Escherichia coli* abundance increases as a result of VR treatment, as it was only upregulated 2 weeks after withdrawal (Figure 6F). This difference might be due to the different initial microbiome state in this experiment, which was performed with distinct groups of animals. In support of this hypothesis, it was shown that the initial state of the intestinal microbiome influences the effect evoked by antibiotics [53]. Thus, the association of *Escherichia coli* outgrowth with the behavioral phenotype is inconsistent and context-dependent, while *Akkermansia muciniphila* up-regulation followed VR treatment in both experiments and was robustly associated with social behavior.

This finding demonstrate that antibiotic combinations can produce unique microbial ecosystem, which can be used as a tool to study the effect of distinct microbial species. Here we used this tool to investigate the correlation between *Akkermansia muciniphila* and other species abundance on female preference that we have previously noted in a transgenic mouse model of colitis [29]. Our experiment on *Akkermansia muciniphila* gavage provides evidence for its influence on social behavior, supporting the hypothesis that mucin-degrading bacteria may play an outstanding role in gut-brain communication [10].

At the same time, the VR withdrawal experiment demonstrates that *Akkermansia muciniphila’s* propagation itself is not an ultimate prerequisite to the impairment of female-related social behavior in males. While female odor preference and female contact time normalized by the 25^th^ day of withdrawal (Figure 5 F-G), *Akkermansia muciniphila* was still significantly upregulated at this point (Figure 6F). This finding is consistent with previous reports demonstrating long-term microbiome changes after antibiotic treatment and withdrawal [54], [55]. Apparently, other bacterial species contribute to the behavioral recovery 25 days after VR withdrawal. Of all identified bacterial species, only *Pseudomonas azotoformans* was both significantly affected by the treatment and proliferated only by the day 25 of withdrawal (Figure 6F). However, this bacterium is largely undescribed in relation to the host-microbiome environment, and it might be difficult to propose its involvement in the regulation of social behaviors.

Alternatively, a combination of microbial taxa could interfere with the outcome of *Akkermansia muciniphila* upregulation and affect metabolism and behavior. It has been shown that *Akkermansia muciniphila* is a potent source of short-chain fatty acids. It primarily produces acetate, and under certain conditions can also synthetize propionate [56], [57]. At the same time, the onset of social disinterest for females following antibiotic treatment aligns with the depletion of *Lachnospiraceae* potentially resulting in the loss of butyrate production (Figure 6F) [58]. In turn, butyrate is a metabolite crucial for neuroplasticity and social behavior through histone deacetylase inhibition and Brain-Derived Neurotrophic Factor regulation [59]. Thus, the microbial community resulting from VR treatment could promote substantial imbalance in key short-chain fatty acids composition. Notably, the social communication phenotype was reconstituted by administering an acetate-propionate mixture to antibiotic-naive mice, suggesting that short-chain fatty acids imbalance itself ‒ rather than specific bacterial taxa – could drive the observed behavioral changes. While *Akkermansia muciniphila*-derived acetate has been shown to support barrier function [60], our findings demonstrate that excessive non-butyrate SCFAs disrupt the expected social preference (Figure 4K-L). The delayed behavioral recovery at day 25 coincides with the upregulation of *Blautia* ‒ known butyrate producer. It implies that even partial restoration of butyrogenic capacity could compensate for acetate/propionate effects. Our data is consistent with studies showing that excessive non-butyrate short-chain fatty acids disrupt dopaminergic signaling in reward circuits and impair social recognition [61], [62]. This supports an emerging paradigm where short-chain fatty acids ratios, rather than absolute concentrations, modulate gut-brain signaling. The persistence of *Akkermansia muciniphila* during behavioral recovery further suggests its role as a metabolic modulator whose effects depend on the broader microbial context. However, this hypothesis is purely theoretical and requires further experimental investigation.

Our study also raises important translational considerations regarding the use of vancomycin and rifampicin. The persistence of behavioral effects long after antibiotic withdrawal suggests potential clinical relevance for understanding long-term consequences of antibiotic use, particularly in vulnerable populations [13].

Finally, the VR-treatment paradigm establishes a novel experimental model for studying microbiome-mediated disruptions in sex-specific social preference behaviors in mice. Our comprehensive behavioral characterization includes validated tests for social interaction (resident-intruder and two-intruder paradigms), olfactory discrimination (odor preference and recognition tests), coupled with multi-omics analyses of gut microbiota composition and brain neurotransmitter metabolism. We propose that this model could become a valuable tool to study sexually dimorphic behaviors and the underlying neural mechanisms in mice.

## 5. Conclusions

In conclusion, this work provides compelling evidence that specific antibiotic-induced microbiota alterations can lead to persistent and selective impairments in social behavior, mediated partially through the brain energy metabolism. These findings broaden our understanding of gut-brain communication and suggest new avenues for investigating and potentially treating social behavior disorders. Future research should focus on elucidating the precise molecular mechanisms linking microbial and metabolic changes to neural function. The finding that *Akkermansia muciniphila* and short-chain fatty acids can adversely modulate social behavior highlights new aspects of the gut-brain axis research. However, these results also demonstrate that potential therapeutic application of *Akkermansia muciniphila* as a probiotic demands further investigation into its context-dependent effects on neural circuits and behavior.

## Author Contributions

Conceptualization, E.K.; methodology, M.M. and E.K.; software, J.A., K.A., and E.K.; validation, M.M., K.A. and J.A.; formal analysis, M.M., K.A., J.A, O.S., Y.T.; investigation, M.M., K.A., J.P, E.A, O.S., Y.T ; resources, E.N.; data curation, E.N.; writing ‒ original draft preparation, E.N.; writing ‒ review and editing, M.M., O.S., J.A., J.P., K.A., Y.T., and E.K.; visualization, J.A. and J.P.; supervision, E.N. and Y.T.; project administration, E.N.; funding acquisition, E.N. All authors have read and agreed to the published version of the manuscript.

## Institutional Review Board Statement

The animal study protocol was approved by the Institutional Ethics Committee of The Institute of Molecular and Cellular Biology of 06.10.2021.

## Data Availability Statement

Metagenomic data can be found here: https://www.ncbi.nlm.nih.gov/bioproject/PRJNA1248659. Metabolomic data is available upon request and will be uploaded on the open resource by the time of publication.

## Acknowledgments

O.S. and Y.T. thank the Ministry of Science and Higher Education of the Russian Federation for granting access to the equipment. The authors thank Dr. E.N. Pavlova for the assistance with monoamine measurements, K.S. Pavlov and Dr. L.V. Boldyreva for technical support with behavioral tests and sample management. We acknowledge the use of generative AI during the final editing phase to improve the manuscript’s readability and language.

